# A Practical Covariance-Based Method for Efficient Detection of Protein-Protein Attractive and Repulsive Interactions in Molecular Dynamics Simulations

**DOI:** 10.1101/2025.03.24.644990

**Authors:** Mert Golcuk, Mert Gur

## Abstract

Molecular dynamics simulations of large protein–protein complexes require scalable analysis. We present a correlation-based workflow that systematically identifies both attractive (stabilizing) interactions, such as salt bridges, hydrogen bonds, and hydrophobic contacts, as well as repulsive (destabilizing) interactions across extensive interfaces. By constructing an inter-protein covariance matrix, filtering residue pairs based on distance, and identifying interactions underlying these correlations, our method focuses computational resources on the most relevant regions of the interface while preserving a high level of detail.

## INTRODUCTION

As both hardware and software technologies continuously advance, the size of protein systems and the length of all-atom molecular dynamics (MD) simulations that can be performed are expanding dramatically. A decade ago, typical simulations involved tens of thousands of atoms and spanned only a few tens of nanoseconds; today, it is not uncommon to simulate hundreds of thousands (or even millions^1-3^) of atoms over microsecond timescales. Consequently, identifying interactions at protein–protein interfaces was once manageable through simple visual inspection or basic scripts. However, modern system sizes and simulation lengths demand more efficient methods to identify and characterize interactions across large protein–protein interfaces. Examples include interactions at oligomerization surfaces of homomers or heteromers, antibody–antigen complexes, motor proteins binding to microtubules, and microtubule-associated proteins (MAPs) binding to microtubules. For instance, the nanobody H11-H4 that binds the SARS-CoV-2 Spike protein has a binding interface spanning approximately 78 residues, while kinesin and dynein each have microtubule interaction surfaces comprising at least 101 residues.

Although a variety of publicly available codes and in-house tools exist to evaluate specific protein– protein interactions, they often rely on criteria such as distance and angle cutoffs for hydrogen bonds^4^ or distance-based cutoffs for salt bridge determination^5^. However, such criteria-based methods can become computationally expensive and challenging to scale to larger systems and longer simulations, as the number of pairwise calculations required to identify interacting residues increases significantly with the size of the system and the duration of the simulation. Furthermore, identifying hydrophobic interactions through simple distance cutoffs is not straightforward. This is because the presence of a water molecule between two hydrophobic residues can disrupt hydrophobic interaction. Additionally, detecting repulsive interactions (i.e., destabilizing interactions), such as unfavorable electrostatic contacts between like-charged residues is complex, since most criteria-based approaches focus primarily on attractive interactions (i.e., stabilizing interactions) and lack specific metrics to systematically identify or quantify these repulsive interactions.

Here, we introduce a new method that integrates and extends our existing approach^6^, in which we demonstrated that constructing a covariance matrix from MD simulations can be used to efficiently identify both the presence and strength of interactions between two bound peptides from MD simulations. The basic idea is that the elements of the covariance matrix indicate how strongly the motions of two residues are correlated and whether these motions occur in the same or opposite directions, thereby revealing potential direct interactions across the interface as well as signatures of allosteric coupling. Unlike the prior covariance analysis, which was applied to a peptide–peptide system, the present workflow introduces a spatial filtering step to specifically select correlated residue pairs that are in contact, as well as an interaction classification step, thus enabling its application to large protein interfaces and the detection of repulsive like-charge interactions. This results in a powerful, efficient, and user-friendly new approach for identifying both attractive interactions (e.g., salt bridges, hydrogen bonds, hydrophobic interactions) and repulsive interactions (e.g., repulsion between like-charged residues) from MD simulations. We demonstrate the effectiveness of this covariance-based workflow by successfully identifying key attractive and repulsive interactions in MD simulations of complexes involving the nanobody H11-H4 bound to multiple SARS-CoV-2 Spike protein variants, including Alpha, Beta, and Omicron.

## METHODS

### Algorithm

The details and implementation of the introduced methodology, comprising three main steps, are described as follows:

### 1. Constructing the Inter-Protein Covariance Matrix

For a protein–protein complex, the inter-protein covariance matrix (**C**)^7^ is computed as:

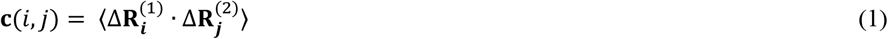

Here 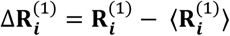 and 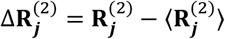 represent the three-dimensional displacement vectors of atom *i* in protein 1 and atom *j* in protein 2, respectively. 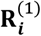and 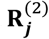denote the position vectors for a given protein conformation (time frame) from an MD simulation, while 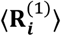 and 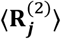 represent the corresponding trajectory-averaged position vectors. We adopt a coarse-grained model based on the C_α_ atom representation. *n* and *m* represent the number of residues considered in *protein 1* and *protein 2* (either all C_α_ of the full sequences or selected subsets), respectively; resulting in a covariance matrix having dimensions of *n* x *m*. However, the model can also be extended to an all-atom representation or to another atom selection, such as backbone atoms.

The normalized covariance (i.e. cross correlation)^7^ is calculated as,

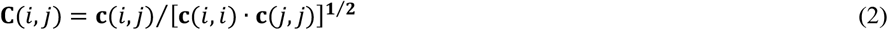

Because of the normalization, the value **C**(*i, j*) is bounded within the range −1 ≤ **C**(*i, j*) ≤ 1. A **C**(*i, j*) value of 1 signifies completely correlated motions between atoms *i* and *j*, while −1 indicates completely anticorrelated motions between these atoms. **C**(*i, j*) values near zero indicate that the motions of the two atoms/residues are essentially uncorrelated.

### 2. Applying a Distance Cutoff to Identify Close Contact Correlations

As a next step, a spatial cutoff is applied to identify residue pairs that come within a defined “interaction distance.” For simplicity, the trajectory-averaged coordinates 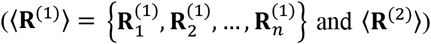, already computed in the previous step, are used. Distance thresholds of 11 Å for positive correlations and 13 Å for negative correlations were selected to align with typical MD simulation parameters. Specifically, a non-bonded interaction cutoff of 12 Å with a switching function starting from 10 Å was applied in the MD simulations. Thus, the chosen thresholds ensure capturing relevant attractive interactions while also accounting for slightly longer-range repulsive correlations between like-charged residues. Any elements of the covariance matrix corresponding to residue pairs whose average inter-residue distance falls outside the cutoff are set to zero and a new ‘close-contact covariance matrix’ is constructed.

Conceptually, when two residues interact, they tend to maintain a preferred equilibrium distance, moving in concert: if one residue moves in a positive direction, the other typically moves similarly, and vice versa. In the covariance matrix, such a concerted motion between residues i and j leads to a positive ensemble average (i.e., a positive element (i,j) of the covariance matrix, 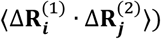. If there is no interaction, the residues move independently, resulting in near-zero 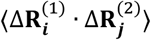. Conversely, if two residues strongly repel each other, they tend to move in opposite directions, resulting in a negative 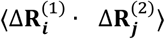 A persistent negative covariance element is observed when a like-charged pair is geometrically confined within the interface by neighboring attractive contacts and by covalent linkage to their respective backbones; these constraints prevent the residues from diffusing far apart, so their repeated ‘push–pull’ motion yields a stable anti-correlation.

### 3. Performing Interaction Analysis

In this step, either 3A or 3B, or preferably both, can be performed to obtain more detailed insights into the protein–protein interactions.

#### 3A. Focused Interaction Calculations

For each close-contact, correlated residue pair identified in the previous step, we assessed the types and frequencies of specific interactions as follows. Salt bridges were identified as interactions between basic nitrogen and acidic oxygen atoms within 6 Å.^5^ Hydrogen bonds were defined as those in which the donor–acceptor distance was ≤3.5 Å and the donor–hydrogen–acceptor angle was ≤30°.^4^ Hydrophobic interactions were characterized by contacts between side-chain carbon atoms within an 8 Å cutoff. Finally, negative correlations were assigned to repulsive interactions that occurred when (i) two basic nitrogen atoms or two acidic oxygen atoms were separated by ≤ 8 Å, or (ii) an aliphatic hydrophobic residue was located within 8 Å of a charged or polar residue.

#### 3B. Interaction Analysis through Visualization

The correlated, close-contact residue pairs identified in Step 3 are visualized on the protein structures using a molecular visualization program (e.g., VMD^8^) to identify those involved in either attractive or repulsive interactions.

Our method, as presented, operates on a single trajectory that predominantly samples conformational fluctuations around one dominant binding mode. If a system populates several markedly different interfacial states (e.g., a protein complex that binds in two alternate orientations or a highly flexible interface that switches configurations), two practical strategies can be followed: (i) analyze each state separately by clustering the trajectory, so that state-specific covariance matrices and interaction maps can be compared directly; or (ii) combine all conformations into a single ensemble, recognizing that the resulting covariance matrix will be a population-weighted average in which contacts unique to an individual state may be attenuated, whereas robust interactions common to all states will still stand out.

## RESULTS AND DISCUSSION

Our new methodology consists of three straightforward steps, demonstrated using an MD trajectory of a protein–protein complex formed by the nanobody H11-H4 and the SARS-CoV-2 Omicron variant Spike protein. First, an inter-protein covariance matrix is generated using the C_α_atoms from the MD trajectory (Fig. 1 step I). The covariance matrix used can be either the unnormalized covariance matrix (eq. 1) or the normalized covariance matrix (eq. 2); in this work, we used the normalized covariance matrix. Next, a distance cutoff is applied based on the average atomic coordinates used during the covariance matrix construction, setting all matrix elements beyond this cutoff to zero and thereby yielding a “close-contact” covariance matrix. This filtering step highlights residue pairs that are both spatially proximal and dynamically coupled, as illustrated in Fig. 1 step II, where a cutoff of 11 Å is applied for positively correlated pairs and 13 Å for negatively correlated pairs. Notably, the close-contact covariance adds dynamic information beyond static distance alone; it ensures the retained pairs exhibit concerted motion indicative of a real interaction, rather than just accidental proximity. In essence, the covariance matrix acts as a filter of interaction significance.

**Figure 1.**
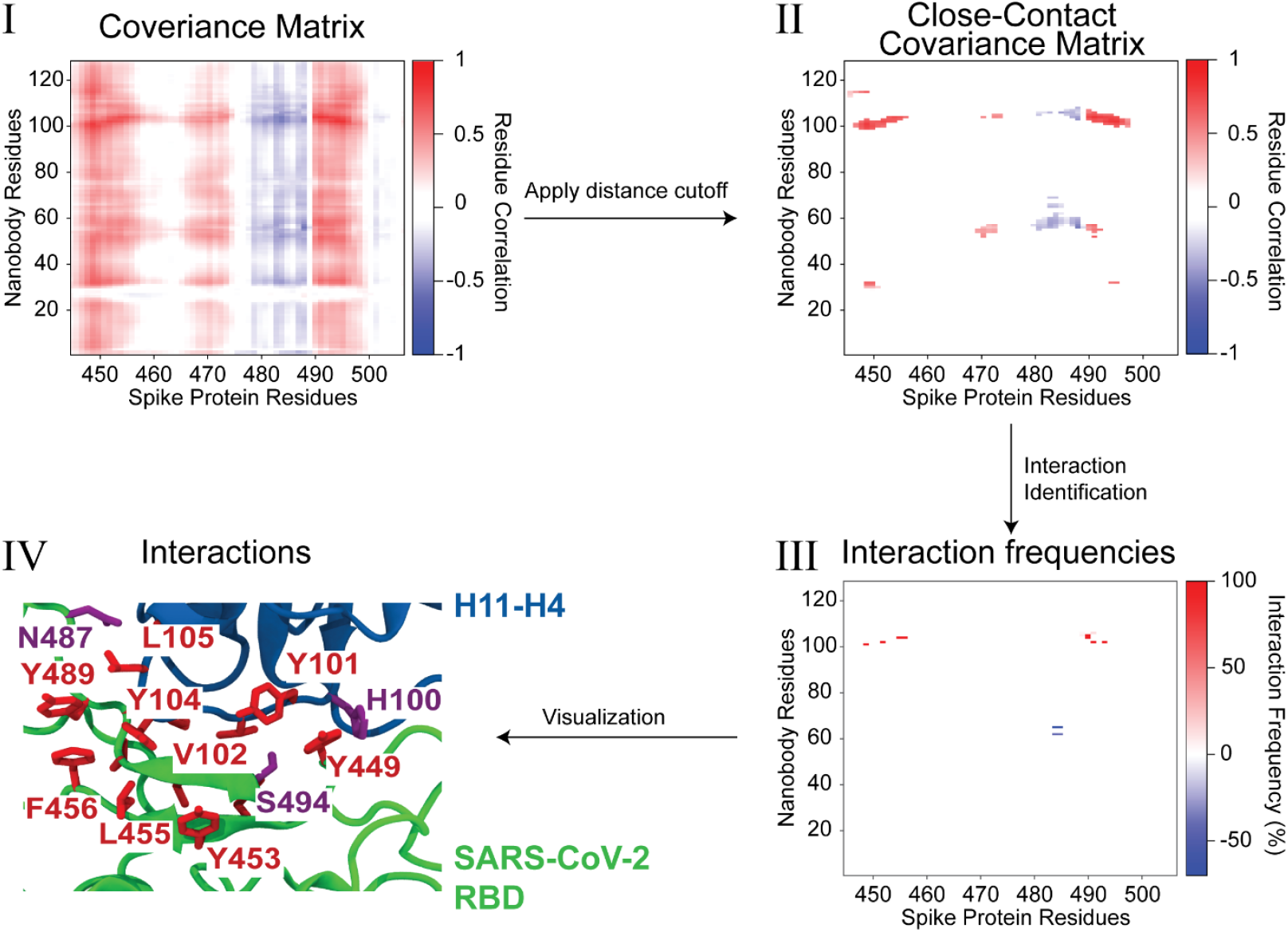
Methodology overview illustrated using the nanobody H11-H4/SARS-CoV-2 Omicron Variant Spike protein complex. **(Step I)** Inter-protein normalized covariance matrix calculated using C_α_ atoms between nanobody H11-H4 residues and SARS-CoV-2 Spike protein residues. Red and blue colors indicate positive and negative covariance elements (residue correlations), respectively, suggesting attractive or repulsive interactions. **(Step II)** Close-contact covariance matrix obtained after applying a spatial distance cutoff (11 Å for positively correlated pairs, 13 Å for negatively correlated pairs), highlighting residue pairs that are spatially proximal and dynamically coupled. **(Step III)** Interaction frequency map displaying the percentage occurrence of interactions for residue pairs identified from the close-contact covariance matrix. All interaction types considered (hydrogen bonds, salt bridges, hydrophobic contacts, and electrostatic repulsions) are shown on a single map. **(Step IV)** Structural visualization of identified interacting residues on nanobody H11-H4 (blue) and the SARS-CoV-2 Spike receptor-binding domain (green). Residues involved in attractive and repulsive interactions are explicitly labeled and colored by physicochemical properties: hydrophobic (red), positively charged (blue), negatively charged (orange), and polar (magenta), providing spatial context to interaction data.

The positive and negative values in the close-contact covariance matrix indicate possible attractive and repulsive interactions, respectively, which will be evaluated in the next step. Using only the residue pairs that pass the close-contact filter, we scan each pair for specific interaction types and, when an interaction is detected, quantify its frequency across the trajectory. Attractive categories include salt bridges, hydrogen bonds, and hydrophobic contacts, whereas repulsive categories encompass same-charge electrostatic repulsion and clashes between hydrophobic and charged residues. During this interaction-classification step, interaction-specific geometric or energetic cutoffs (see Methodology) replace the initial 11 Å/13 Å distance threshold. Restricting the analysis to these pre-selected correlated pairs markedly reduces computational cost by focusing on residues most likely to interact. Fig. 1 step III illustrates the resulting interaction frequency map between the nanobody H11-H4 and the SARS-CoV-2 Spike protein after these three steps. For brevity, all interaction types are shown combined into a single frequency map. However, interactions could alternatively be presented separately as multiple matrices, with individual maps for salt bridges, hydrogen bonds, hydrophobic contacts, and electrostatic repulsions. Fig. 1 visually shows the positions of the correlated close-contact residues displayed on the structures of the nanobody H11-H4 and the SARS-CoV-2 Spike protein. By following this workflow, researchers can streamline protein–protein interaction analyses, reduce data complexity, and achieve clearer insights that can guide subsequent computational and experimental investigations.

To further evaluate the accuracy and effectiveness of our methodology, we applied it to MD simulation trajectories (400 ns length each) from our previous studies^9, 10^ investigating interactions between the nanobody H11-H4 and multiple SARS-CoV-2 Spike protein variants, including Alpha, Beta, and Omicron. Our results demonstrate that our proposed method reliably and efficiently identifies key interactions, consistent with those found using conventional, more time-consuming interaction analysis workflows, in which all possible residue-residue interactions are exhaustively evaluated (Fig. 2). For the WT as well as the Alpha variant, no residue pair exhibited a strong negative covariance element within the 13 Å cutoff, hence the close-contact matrix in Fig. 1 contains only positive (red) values. Furthermore, the overall shape of the interaction map based on the covariance matrix shows highly similar patterns, highlighting the similarity between binding poses of nanobody H11-H4 to the WT and Alpha variant. By contrast, several new anticorrelated residue pairs emerge in the Beta and Omicron variants. These anticorrelations can be traced to repulsive interactions between residues R52–G56 in the CDR2 loop and T99–Y116 in the CDR3 loop of H11-H4. In the Beta variant, the E484K substitution breaks the stabilizing E484–R52 salt bridge and, because E → K reverses the charge, introduces electrostatic repulsion between the RBD and R52 of the nanobody. In the Omicron variant, the E484A substitution likewise abolishes the salt bridge, eliminating the favorable electrostatic attraction altogether. Additional Omicron-specific substitutions further destabilize several hydrophobic and hydrogen-bond contacts mediated by the CDR2 and CDR3 loops of H11-H4.

**Figure 2.**
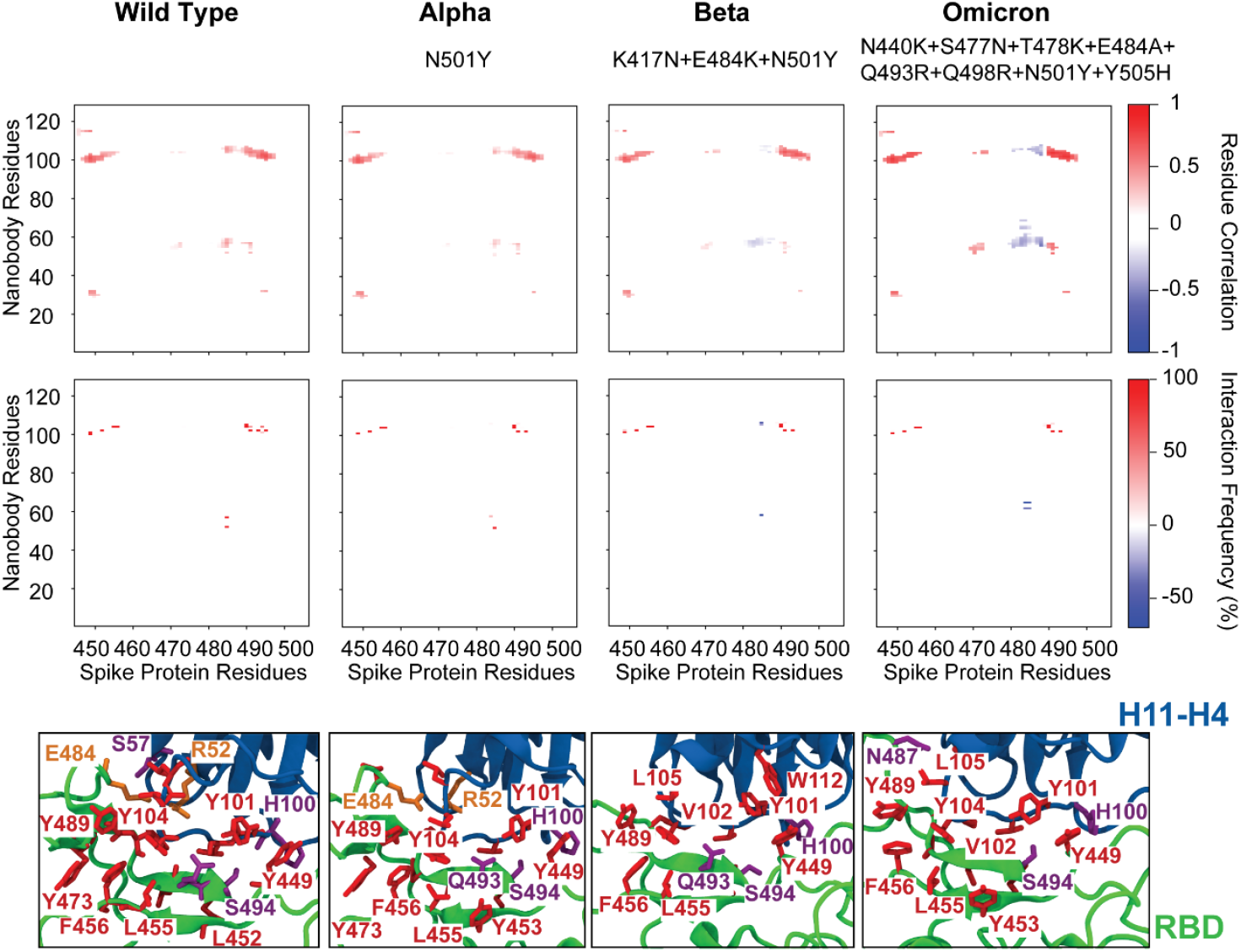
Comparison of nanobody H11-H4 interactions with wild-type and variant SARS-CoV-2 Spike proteins. Top panels display close-contact covariance matrices illustrating residue correlations between nanobody H11-H4 and SARS-CoV-2 Spike receptor-binding domain (RBD) for WT, Alpha, Beta, and Omicron variants. Variant-specific RBD mutations are indicated on each panel, e.g., N501Y in Alpha). Positive correlations (red) in the close-contact covariance matrices indicate attractive interactions, whereas negative correlations (blue) represent repulsive interactions. Middle panels show interaction frequency maps corresponding to each variant, indicating the percentage occurrence of specific residue-residue interactions. Bottom panels visualize interactions on protein structures, highlighting residues involved in attractive and repulsive interactions. Residues are explicitly labeled and colored by physicochemical properties: hydrophobic (red), positively charged (blue), negatively charged (orange), and polar (magenta). Differences in interaction patterns between variants illustrate the impact of Spike protein mutations on nanobody binding.

For the H11-H4/Spike complex (whether WT or mutant), all major contacts (e.g., hydrogen bonds and salt bridges) identified by an exhaustive frame-by-frame search were also captured by our covariance-based workflow, with no significant interactions missed. Notably, the covariance approach dramatically reduced the number of required calculations. An exhaustive brute-force method would require inter-residue distance calculations on approximately 500 residue pairs, whereas our covariance-based filtering reduced this step to ∼80 residue pairs, roughly an order of magnitude reduction. Since distance calculations are computationally expensive and must be computed for every sampled conformation of the trajectory, reducing the number of evaluated residue pairs significantly decreases the computational load. For example, in our study we analyzed 8,000 conformations for each of the 400 ns long simulation; thus, every additional residue pair evaluated would have added 8,000 distance calculations to the computational workload. Our covariance-based method therefore achieves comparable results to brute-force analysis but with substantially improved computational efficiency.

## CONCLUSION

We introduced a covariance-based method that efficiently identifies attractive and repulsive interactions at protein–protein interfaces from MD simulations. In addition to simplifying interaction identification, our method offers a notable advantage: traditional approaches often treat interactions as binary (present or absent based on cutoffs) and generally do not reveal the spatial extent of an interaction’s influence. In contrast, the correlation (covariance) based approach presented here captures not only the presence of interactions but also the dynamic coupling between residues. This enables insight into how specific attractive or repulsive interactions propagate through the protein interface and influence the local binding environment, an aspect that standard interaction analysis methods typically overlook. Furthermore, our covariance-based approach can readily be extended to analyze more complex scenarios, such as interactions within triple or multiprotein systems, facilitating deeper understanding of larger biomolecular assemblies. Constructing very large covariance matrices can become memory-intensive for large protein complexes if an all-atom resolution is used instead of the coarse-grained approach described here. This can be mitigated by pre-filtering residue pairs based on a static structure, such as the trajectory averaged structure or a representative MD conformation corresponding to the most frequently sampled state in the simulation. This approach not only decreases memory usage but also reduces computational load by calculating covariance only for residue pairs that meet the pre-filtering distance criterion. All identified interactions exhibited corresponding normalized covariance values either below -0.08 or above 0.1, clearly indicating that non-zero covariance elements strongly suggest the presence of an interaction. Notably, non-zero covariance elements frequently appear for residue pairs lying beyond the spatial cutoff owing to indirect dynamical coupling. For example, if residue *i* moves in concert with residue k and residue k moves oppositely to residue j, the transitive coupling produces a negative covariance element for the distant i*–*j pair; if the intermediate couplings share the same sign, a positive element results. Such long-range covariance patterns are widely interpreted as hallmarks of allosteric communication pathways that relay motions through the protein framework.^11^ Our distance filter therefore serves a pragmatic purpose, focusing the detailed interaction analysis on physically proximate pairs, while the full covariance map can still be exploited to explore putative allosteric networks.

## Abbreviations

C_α_: Carbon alpha
MD: Molecular dynamics
RBD: Receptor binding domain
WT: Wild type
MAPs: Microtubule-associated proteins

## Data and Software Availability

All data and code developed for this study have been deposited on GitHub.github.com/golcukm/Covariance_PPI.

